# MicroLive: An Image Processing Toolkit for Quantifying Live-cell Single-Molecule Microscopy

**DOI:** 10.1101/2025.09.25.678587

**Authors:** Luis U. Aguilera, William S. Raymond, Rhiannon M. Sears, Nathan L. Nowling, Brian Munsky, Ning Zhao

**Affiliations:** Department of Biochemistry and Molecular Genetics, University of Colorado-Anschutz Medical Campus, 80045, CO, USA; School of Biomedical and Chemical Engineering, Colorado State University, 80523, CO, USA

**Keywords:** image processing, live-cell imaging, single-molecule tracking, translation, transcription, autocorrelation function, colocalization

## Abstract

Advances in live-cell fluorescence microscopy have enabled us to visualize single molecules (such as mRNAs and nascent proteins) in real time with high spatiotemporal resolution. However, these experiments generate large datasets that require complex computational processing pipelines to derive meaningful and quantitative information, which is a technical barrier for many researchers. To address this barrier, here, we introduce `MicroLive`, an open-source Python-based application for quantifying live-cell microscopy images. `MicroLive` provides an interactive Graphical User Interface (GUI) to perform key tasks, including cell segmentation, photo-bleaching correction, single-particle detection/tracking, spot intensity quantification, inter-channel colocalization, and time-series correlation analysis. As a ground-truth testing dataset, we used synthetic live-cell imaging data generated with the rSNAPed toolkit, demonstrating accurate extraction of biologically relevant parameters. Microscopy images of U-2 OS cells expressing a gene construct smHA-KDM5B-BoxB-MS2 were used to demonstrate the use of this software.

**Availability and implementation:** `MicroLive` is distributed under a GPLv3 license and available on GitHub. https://github.com/ningzhaoAnschutz/microlive.

## 1 Introduction

Live-cell single-molecule microscopy is transforming our understanding of gene expression (1). Novel imaging strategies combine advanced live-cell fluorescence microscopy and genetically encoded fluorescent tagging systems, such as intrabody systems (e.g., SunTag (2; 3) and frankenbodies (4; 5)) and RNA stem-loop systems (e.g., MS2 (6) and PP7 stem loops (7)), allowing researchers to visualize nascent proteins and mRNAs directly in real time in live cells (8). These technological advances have revealed complex phenomena, such as single mRNA translation (3; 9; 10; 11; 12), translation bursts (13), frameshifting (14), and IRES-initiated translation (15). However, extracting meaningful information from live-cell microscopy is a complicated multi-step process, including cell segmentation, diffraction-limited fluorescent spots detection, linking detected spots across frames, quantifying spot intensity, and determining colocalization or temporal correlations between imaging channels (16). Performing these analyses often requires a combination of software tools (e.g., ImageJ (17)) and custom scripts, which creates a technical barrier for many researchers. To address this, we introduce **MicroLive**, a user-friendly Graphical User Interface (GUI) platform for live-cell single-molecule imaging analysis. It enables users to load multi-dimensional images and implement a complete image-processing pipeline through a point-and-click graphical interface, as shown in Figure 1.

**Figure 1:**
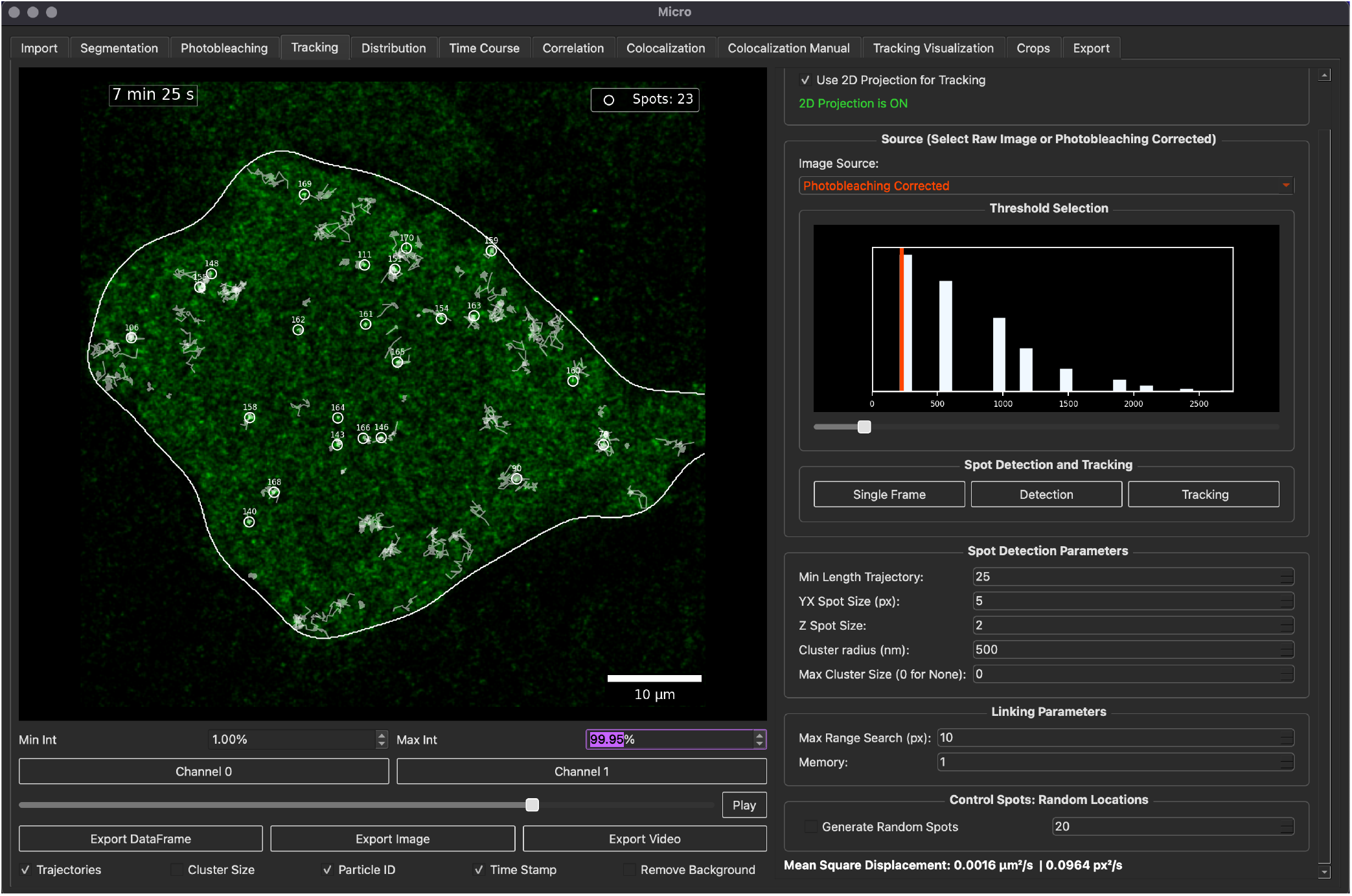
A snapshot of `MicroLive`. Multiple tabs allow users to perform cell segmentation, spot detection, particle tracking, intensity calculation, colocalization, and correlation analysis. The processed image represents a U-2 OS cell expressing the smHA-KDM5B-BoxB-MS2 gene construct.

## 2 Method and Implementation

`MicroLive` is implemented in Python with a modular architecture: the core image processing routines are coded as classes and reside in `src/microscopy.py`, while the code generating the GUI is located in `gui/micro.py`. `MicroLive` applies parallel computing to accelerate compute-intensive tasks, such as processing multiple-frame images. Additionally, the GUI loads on-demand individual 2D frames or 3D stacks from the disk only when the frame or stack is displayed or processed, then frees the memory immediately. This on-demand strategy minimizes memory usage and supports large datasets.

`MicroLive` supports standard microscopy files like multi-dimensional TIFF or LIF. For unstructured TIFF files, a Jupyter Notebook is provided to convert them into the standard format used in the GUI for downstream image analyses. Once loaded in `MicroLive`, microscopy images are internally converted into `uint16` NumPy arrays. As a pre-processing step, the user can apply an exponential decay to correct the loss of fluorescence intensity due to photobleaching (18). Segmentation employs either a manual outline tool that allows users to draw Regions of Interest (ROI) or a watershed method to define ROIs (19). Masks are overlaid on images for validation and used to map detected spots to individual cells.

Particles or spots are detected using the TrackPy library (20) for 2D images or Big-FISH (21) for 3D images. Detected particles can be labeled as clusters if their size is multiple times larger than the user-defined particle size. Users can filter out large clusters that may represent aggregates. Particle trajectories are constructed using a nearest-neighbor algorithm with tunable displacement and memory parameters. Particle linking is supported for both 2D and 3D images. Importantly, the code offers visualization of the linked trajectories and the particle identifier, allowing users to detect and correct tracking errors in real-time. For multi-color images, `MicroLive` allows automated methods to detect colocalized spots by using a convolutional neural network (22) trained on manually-annotated images combined with augmented synthetic data to predict the presence of particles in a given ROI. Automated colocalization can be manually curated to reduce false positives and false negatives. Fluorescence intensities for the detected spots can be extracted over time using multiple methods, including total intensity, Gaussian fits, and background subtraction methods. `MicroLive` computes auto- and cross-correlation functions to reveal kinetic parameters such as intensity fluctuations, dwell times, and inter-signal delays (23). All data generated by the GUI can easily be exported as CSV files for downstream processing. Metadata containing all the parameters and thresholds used during the image processing steps is automatically generated and exported to ensure reproducibility. A complete description of the methods used in `MicroLive` is given in the Supplementary File.

## 3 Validation and Results

To verify our code with a ground truth dataset, we used synthetic movies generated with rSNAPed (24). For this, a synthetic dataset containing 360 frames with a 5-sec frame interval, two color channels, 512 × 512 pixels, and 80 mRNA spots was generated to model mRNA translation using an initiation rate of 0.04 1/sec and an elongation rate of 5 aa/sec. Photobleaching was simulated using a decreasing exponential function with a decay rate of 0.001 1/sec. Ground-truth positions and intensities were retained for benchmarking, and recovered results are provided in Supplementary Figure S2, showing a strong agreement between the tracked particles and the values used for the simulation. For example, `MicroLive` accurately detected more than 70 mRNA spots at all time points. `MicroLive` estimated an intensity decay of 0.001 1/sec. Additionally, to determine whether `MicroLive` can correctly extract temporal intensity from the images, we calculated the autocorrelation function of nascent protein intensity traces and extracted initiation and elongation rates as described by Larson et al. (25). We obtained a de-correlation time of ≈ 365 sec, corresponding to an elongation rate of 5.2 aa/sec, and a value for the autocorrelation function at lag zero *G*(0) = 0.07, corresponding to an initiation rate of 0.038 1/sec. A side-by-side comparison between synthetic data generated with the rSNAPed library and `MicroLive` recovered values is given in the Supplementary File Table S1 and Figure S2.

To test `MicroLive`, we analyzed microscopy translation images of a gene construct smHA-KDM5B-BoxB-MS2 (26) in live U-2 OS cells. The smHA-KDM5B-BoxB-MS2 (plasmid sequence is provided) construct consists of an N-terminal spaghetti monster HA (smHA) tag (27) (including 10x HA tags) fused to our protein-of-interest KDM5B, 15x BoxB (26; 28; 29) and 24x MS2 stem-loops (6) in the 3’ untranslated region (UTR). Translation spots were identified by colocalized nascent protein spots visualized by anti-HA-frankenbody-HaloTag (4) stained with JF646 dyes and mRNA spots labeled by tandem MS2 Coat Protein (tdMCP) (6) fused with tandem monomeric StayGold (tdmSG) (30; 31). The mRNA spots were tethered to the plasma membrane through the interaction between *λ*N-CAAX and the BoxB stem-loops (29). The translation images were collected using a Leica Stellaris 5 confocal microscope with a 63x oil objective for 600 frames at 1 frame per second (fps) rate with a single *z* plane. Images were loaded into the GUI to perform cell segmentation, spot detection, intensity calculation, and correlation analyses. The complete quantification is presented in the Supplementary File.

## 4 Conclusion

`MicroLive` is a toolbox to quantify single-molecule microscopy images. `MicroLive` unifies commonly used image processing tasks such as cell segmentation, spot detection, particle tracking, co-localization, and correlation analyses within a single accessible platform. The user-friendly GUI platform lowers the barrier for non-programmers to perform complex image-processing tasks. `MicroLive` is open-source with a GPLv3 license.

## Acknowledgments

Plasmid smHA-KDM5B-BoxB-MS2 was kindly provided by Gabriel Galindo in the Stasevich lab at Colorado State University.

## References

[1] Tatsuya Morisaki, O’Neil Wiggan, and Timothy J. Stasevich. Translation dynamics of single mRNAs in live cells. Annual Review of Biophysics. 53(2024), pages 65–85. 2024.

[2] Marvin E. Tanenbaum, Luke A. Gilbert, Lei S. Qi, Jonathan S. Weissman, and Ronald D. Vale. A protein-tagging system for signal amplification in gene expression and fluorescence imaging. Cell., 159(3), pages 635–646. 2014.

[3] Xiaowei Yan, Tim A. Hoek, Ronald D. Vale, and Marvin E. Tanenbaum. Dynamics of translation of single mRNA molecules in vivo. Cell. 165(4), pages 976–989. 2016.

[4] Ning Zhao, Kouta Kamijo, Philip D. Fox, Haruka Oda, Tatsuya Morisaki, Yuko Sato, Hiroshi Kimura, and Timothy J. Stasevich. A genetically encoded probe for imaging nascent and mature HA-tagged proteins in vivo. Nature Communications. 10(1). pages 2947. 2019.

[5] Yang Liu, Ning Zhao, Masato T. Kanemaki, Yotaro Yamamoto, Yoshifusa Sadamura, Yuma Ito, Makio Tokunaga, Timothy J. Stasevich, and Hiroshi Kimura. Visualizing looping of two endogenous genomic loci using synthetic zinc-finger proteins with anti-FLAG and anti-HA frankenbodies in living cells. Genes to Cells. 26(11), pages 905–926. 2021.

[6] Edouard Bertrand, Pascal Chartrand, Matthias Schaefer, Shailesh M. Shenoy, Robert H. Singer, and Roy M. Long. Localization of ASH1 mRNA particles in living yeast. Molecular Cell. 2(4), pages 437–445. 1998.

[7] Jeffrey A. Chao, Yury Patskovsky, Steven C. Almo, and Robert H. Singer. Structural basis for the coevolution of a viral RNA–protein complex. Nature Structural & Molecular Biology. 15(1). pages 103–105. 2008.

[8] Rhiannon M Sears, Nathan L Nowling, Jake Yarbro, and Ning Zhao. Expanding the tagging toolbox for visualizing translation live. Biochemical Journal. 482(3), pages 147–165. 2025.

[9] Tatsuya Morisaki, Kenneth Lyon, Keith F. DeLuca, Jennifer G. DeLuca, Brian P. English, Zhengjian Zhang, Luke D. Lavis, Jonathan B. Grimm, Sarada Viswanathan, Loren L. Looger, Timothee Lionnet, and Timothy J. Stasevich. Real-time quantification of single RNA translation dynamics in living cells. Science. 352(6292), pages 1425–1429. 2016.

[10] Chong Wang, Boran Han, Ruobo Zhou, and Xiaowei Zhuang. Real-time imaging of translation on single mRNA transcripts in live cells. Cell. 165(4), pages 990–1001. 2016.

[11] Bin Wu, Carolina Eliscovich, Young J. Yoon, and Robert H. Singer. Translation dynamics of single mRNAs in live cells and neurons. Science. 352(6292), pages 1430–1435. 2016.

[12] Xavier Pichon, Amandine Bastide, Adham Safieddine, Racha Chouaib, Aubin Samacoits, Eugenia Basyuk, Marion Peter, Florian Mueller, and Edouard Bertrand. Visualization of single endogenous polysomes reveals the dynamics of translation in live human cells. Journal of Cell Biology. 214(6), pages 769–781. 2016

[13] Nathan M. Livingston, Jiwoong Kwon, Oliver Valera, James A. Saba, Niladri K. Sinha, Pranav Reddy, Blake Nelson, Clara Wolfe, Taekjip Ha, Rachel Green, Jian Liu, and Bin Wu. Bursting translation on single mRNAs in live cells. Molecular Cell. 83(13), pages 2276–2289. 2023.

[14] Kenneth Lyon, Luis U. Aguilera, Tatsuya Morisaki, Brian Munsky, and Timothy J. Stasevich. Live-cell single RNA imaging reveals bursts of translational frameshifting. Molecular Cell. 75(1), pages 172–183. 2019.

[15] Amanda Koch, Luis Aguilera, Tatsuya Morisaki, Brian Munsky, and Timothy J. Stasevich. Quantifying the dynamics of IRES and cap translation with single-molecule resolution in live cells. Nature Structural & Molecular Biology. 27(12), pages 1095–1104. 2020.

[16] Deepak Khuperkar, Tim A. Hoek, Stijn Sonneveld, Bram M.P. Verhagen, Sanne Boersma, and Marvin E. Tanenbaum. Quantification of mRNA translation in live cells using single-molecule imaging. Nature Protocols. 15(4), pages 1371–1398. 2020.

[17] Johannes Schindelin, Ignacio Arganda-Carreras, Erwin Frise, Verena Kaynig, Mark Longair, Tobias Pietzsch, Stephan Preibisch, Curtis Rueden, Stephan Saalfeld, Benjamin Schmid, Jean-Yves Tinevez, Daniel James White, Volker Hartenstein, Kevin Eliceiri, Pavel Tomancak, and Albert Cardona Fiji: an open-source platform for biological-image analysis. Nature Methods 9(7) pages 676–682. 2012.

[18] Kota Miura. Bleach correction ImageJ plugin for compensating the photobleaching of time-lapse sequences. F1000Research. 9, pages 1494. 2020.

[19] Luc Vincent, and Pierre Soille. Watersheds in digital spaces: an efficient algorithm based on immersion simulations. IEEE Transactions on Pattern Analysis & Machine Intelligence. 13(6), pages 583–598. 1991.

[20] Daniel B. Allan, Thomas Caswell, Nathan C. Keim, Casper M. van der Wel, and Ruben W. Verweij. Softmatter/trackpy: V0.7. Zenodo 10.5281/zenodo.16089574. 2025.

[21] Arthur Imbert, Wei Ouyang, Adham Safieddine, Emeline Coleno, Christophe Zimmer, Edouard Bertrand, Thomas Walter, and Florian Mueller. FISH-quant v2: a scalable and modular tool for smFISH image analysis. RNA. 28(6), pages 786–795. 2022.

[22] Jay M. Newby, Alison M. Schaefer, Phoebe T. Lee, M. Gregory Forest, and Samuel K. Lai. Convolutional neural networks automate detection for tracking of submicron-scale particles in 2D and 3D. Proceedings of the National Academy of Sciences. 115(36), pages 9026–9031. 2018.

[23] Antoine Coulon, and David R. Larson. Fluctuation analysis: dissecting transcriptional kinetics with signal theory. Methods in Enzymology. Vol. 572, pages 159–191. 2016.

[24] William S. Raymond, Sadaf Ghaffari, Luis U. Aguilera, Eric Ron, Tatsuya Morisaki, Zachary R. Fox, Michael P. May, Timothy J. Stasevich, and Brian Munsky. Using mechanistic models and machine learning to design single-color multiplexed nascent chain tracking experiments. Frontiers in Cell and Developmental Biology. 11, 1151318. 2023.

[25] Daniel R. Larson, Daniel Zenklusen, Bin Wu, Jeffrey A. Chao, and Robert H. Singer. Real-time observation of transcription initiation and elongation on an endogenous yeast gene. Science. 332(6028), pages 475–478. 2011.

[26] Gabriel Galindo, Gretchen M. Fixen, Amelia Heredia, Tatsuya Morisaki, and Timothy J. Stasevich. All Probes Plasmids (APPs) for multicolor and long-term tracking of single-mRNA translation dynamics. Molecular Biology of the Cell. 36(6):mr6. 2025.

[27] Sarada Viswanathan, Megan E Williams, Erik B Bloss, Timothy J Stasevich, Colenso M Speer, Aljoscha Nern, Barret D Pfeiffer, Bryan M Hooks, Wei-Ping Li, Brian P English, Teresa Tian, Gilbert L Henry, John J Macklin, Ronak Patel, Charles R Gerfen, Xiaowei Zhuang, Yalin Wang, Gerald M Rubin, and Loren L Looger High-performance probes for light and electron microscopy. Nature Methods. 12(6), pages 568–576. 2015.

[28] Charlotte A. Cialek, Gabriel Galindo, Tatsuya Morisaki, Ning Zhao, Taiowa A. Montgomery, and Timothy J. Stasevich. Imaging translational control by Argonaute with single-molecule resolution in live cells. Nature Communications. 13(1), pages 3345. 2022.

[29] Manuela Schärpf, Heinrich Sticht, Kristian Schweimer, Markus Boehm, Silke Hoffmann, and Paul Rösch. Antitermination in bacteriophage λ: The structure of the N36 peptide-boxB RNA complex. European Journal of Biochemistry. 267(8), pages 2397–2408. 2000.

[30] Masahiko Hirano, Ryoko Ando, Satoshi Shimozono, Mayu Sugiyama, Noriyo Takeda, Hiroshi Kurokawa, Ryusaku Deguchi, Kazuki Endo, Kei Haga, Reiko Takai-Todaka, Shunsuke Inaura, Yuta Matsumura, Hiroshi Hama, Yasushi Okada, Takahiro Fujiwara, Takuya Morimoto, Kazuhiko Katayama, and Atsushi Miyawaki A highly photostable and bright green fluorescent protein. Nature Biotechnology. 40(7), pages 1132–1142. 2022.

[31] Ryoko Ando, Satoshi Shimozono, Hideo Ago, Masatoshi Takagi, Mayu Sugiyama, Hiroshi Kurokawa, Masahiko Hirano, Yusuke Niino, Go Ueno, Fumiyoshi Ishidate, Takahiro Fujiwara, Yasushi Okada, Masaki Yamamoto, and Atsushi Miyawaki StayGold variants for molecular fusion and membrane-targeting applications. Nature Methods. 21(4), pages 648–656. 2024.

