## Supplemental Information for "MicroLive: An Image Processing Toolkit for Quantifying Live-cell Single-Molecule Microscopy"

#### General Architecture

`MicroLive` is a Python library offering a Graphical User Interface (GUI) for quantitative analysis of live-cell single-molecule microscopy images. As illustrated in Figure S1, the GUI is organized into five sequential modules (shown in green in Figure S1), each of which performs a distinct major function: i) **Utilities** for data import and export, ii) **Preprocessing** for segmentation and photobleaching correction, iii) **Detection** for particle tracking, time course, and colocalization, iv) **Statistical** for distributions and correlation analysis, and v) **Visualization** for displaying crops and trajectories. Under the five modules, there are 12 tabs shown in purple in Figure S1, and ordered from left to right in the GUI (Figure 1), reflecting the natural progression of an imaging analysis workflow.

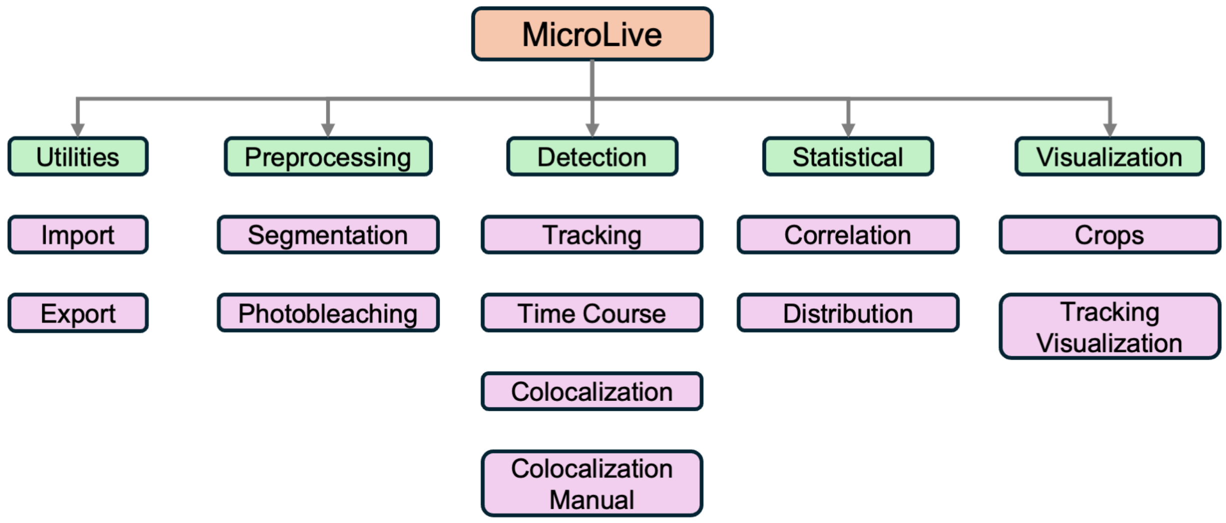


Figure S1: Code Architecture of `MicroLive`. The diagram shows the main code functional modules and tabs.

#### Image Loading

Images can be imported into the `MicroLive` GUI via the **Open File** button located in the **Import** tab. `MicroLive` supports standard microscopy files, such as multi-dimensional TIFF/OME-TIFF and LIF. Once loaded, the GUI automatically extracts critical metadata including spatial dimension, pixel dimension, frame time intervals, laser intensities, and channel configuration. If any critical metadata is missing, the GUI prompts the user to input values manually to ensure accurate unit conversion. For unstructured TIFF images, a Jupyter Notebook (`notebooks/converter.ipynb`) is provided to convert these images into the standard format. Once images are loaded into `MicroLive`, they are mapped into a standard five-dimensional array format (time, $z$, $y$, $x$, channel), ensuring consistent downstream processing. This process ensures that subsequent quantitative analyses, such as spot intensity measurement and particle tracking, are correctly performed and interpreted in terms of physical units and axis dimensions.

### Cell Segmentation

To define cellular Regions of Interest (ROIs) for analysis, the `MicroLive` GUI supports both manual and automated segmentation methods. For manual segmentation, users can define ROIs by drawing directly on images using a built-in interactive polygon drawing tool. For automated segmentation, a watershed-based algorithm is applied with a user-adjustable slider to define the segmentation threshold (1). Optionally, maximum temporal projections of the movies can be used as input for segmentation, which accounts for cell movement over time. The user-defined ROIs are stored as binary mask arrays to define the regions for all downstream analyses.

### Photobleaching Correction

The **Photobleaching** tab corrects for progressive fluorescence decay by fitting an exponential model to the mean intensity time course and then rescaling each frame based on the fitted decay curve (2). The detailed method is shown as follows:

$$I_{\mathrm{fit}}(t) = I_0\, e^{-kt}, \tag{1}$$

here, $I_{\mathrm{fit}}(t)$ is the fitted fluorescence intensity at time $t$, $I_0$ is the initial intensity, and $k$ is the decay constant.

To correct the raw images for photobleaching in channel $c$, we used a unit-normalized approach that approximates the intensity at $t = 0$ to remove the global photobleaching in all subsequent frames. Specifically, we introduce the following *normalized* correction factor applied to each frame:

$$f^{(c)}(t) = \frac{I_{\mathrm{fit}}^{(c)}(0)}{I_{\mathrm{fit}}^{(c)}(t)}, \tag{2}$$

then each raw image at time $t$ in channel $c$ is multiplied by the *normalized* correction factor, effectively restoring its intensity to the estimated unbleached level, as shown below,

$$I_{\mathrm{corr}}^{(c)}(t) \;=\; f^{(c)}(t)\, I_{\mathrm{raw}}^{(c)}(t). \tag{3}$$

### Spot Properties

#### Spot Intensity

The `MicroLive` GUI provides three methods for quantifying spot intensities.

- The first method uses local background subtraction(3). For this, the mean intensities of the spot region $D$ and an annular region $R$ around the spot (local background) are determined, and then the local background intensity is subtracted from the spot intensity. That is:

$$I_{\mathrm{spot}}(x,y) = \frac{1}{s_{\mathrm{spot}}^2} \sum_{(x,y)\in D} I(x,y) - \frac{1}{s_{\mathrm{bg}}^2 - s_{\mathrm{spot}}^2} \sum_{(x,y)\in R} I(x,y), \tag{4}$$

where $s_{\text{spot}}$ is the user-defined spot size in pixels, and $s_{\text{bg}}$ is defined as an outer region extending 3 pixels beyond $s_{\text{spot}}$ in each $xy$ direction.

- For the second method, the spot intensity is estimated by fitting a 2D Gaussian function to determine its peak amplitude $I_0$ above the background. The intensity profile of a spot is modeled as

$$I_{\text{spot}}(x, y) = I_{\text{bg}} + I_0 \exp\Big(-\tfrac{1}{2}\Big[\frac{(x-x_0)^2}{\sigma_x^2} + \frac{(y-y_0)^2}{\sigma_y^2}\Big]\Big), \tag{5}$$

where $I_{\text{bg}}$ is the local background level; $\sigma_x$, $\sigma_y$ are the fitted spot widths along the $x$ and $y$ axes, respectively; $I_0$ is the peak amplitude of the Gaussian function; $x_0$ and $y_0$ are the coordinates of the center of mass of the detected spot.

- The third method calculates the integrated intensity by summing the pixel intensities for all pixels within the spot region $D$:

$$I_{\text{spot}}(x, y) = \sum_{(x,y)\in D} I(x, y). \tag{6}$$

By default, the three methods are applied automatically by the GUI, and their results are exported into the final dataframe.

#### Spot Size

For each detected spot, the GUI first fits its fluorescence profile to a 2D Gaussian function (see Eq. 5). The spot size is then reported as the Full Width at Half Maximum (FWHM), defined as:

$$\text{FWHM} = 2\sqrt{2\ln 2} \times \sigma_{xy}, \approx 2.355 \times \sigma_{xy}, \tag{7}$$

where $\sigma_{xy}$ represents the mean of $\sigma_x$ and $\sigma_y$ obtained from Eq. 5.

For 3D particle tracking, in cases where large spots are detected (e.g., clusters), the GUI computes a cluster size metric using the Big-FISH library(4). This metric represents the number of individual spot detections grouped within a defined area. Identified clusters can be optionally excluded from downstream analyses.

#### Signal-to-Noise Ratio

The Signal-to-Noise Ratio (SNR) of each spot is calculated by dividing its intensity (calculated by using Eq. 4) by the local background noise (i.e., the standard deviation of the pixel intensity in the region $R$, $\sigma_{\text{bg}}(x, y)$).

$$\text{SNR} = \frac{I_{\text{spot}}(x, y)}{\sigma_{\text{bg}}(x, y)}. \tag{8}$$

### Particle Tracking

The tracking module enables automated detection and linking of particles in either two or three dimensions. There are two main functionality modes: **Detection** and **Tracking**. In the **Detection** mode, the GUI detects all particles in each frame, but no particle linking is applied. The **Tracking** mode implements both particle detection and particle linking. In 2D tracking (the default), $z$ planes in each movie are maximum projected, and the TrackPy library(4) is used for spot detection and frame-to-frame linking of particle trajectories(5). In 3D tracking, `MicroLive` employs the Big-FISH library to detect fluorescent spots in 3D (across all $z$ planes) and applies TrackPy for linking trajectories in x–y–z space over time.

##### Time Courses

Time course analysis can be used to track the temporal evolution of the image properties. For each frame, the number of detected spots in an ROI can be counted to produce a particle count vs. time curve. In addition, average intensity, SNR, spot size, and spot intensity values for all spots in the ROI can be plotted over time. All measurements can be plotted after photobleaching correction to ensure that observed temporal changes reflect true biological dynamics rather than gradual intensity loss by bleaching.

#### Spot Colocalization

When analyzing multi-channel (multi-color) fluorescence images, the GUI enables users to detect spots in a reference channel and assess their colocalization with spots in a test channel. The GUI provides the following three approaches to determine spot colocalization.

- **Machine learning approach**. The `MicroLive` GUI uses a machine learning classifier to determine spot colocalization in an image crop. This classifier is implemented as a Convolutional Neural Network (CNN, `ParticleDetectionCNN` in `src/ML_SpotDetection.py`) trained on a mix of human-annotated real microscopy images and synthetically generated spot images. Its architecture comprises two convolutional blocks `conv1` ($1 \to 32$ channels, $3 \times 3$ kernel, padding=1) and `conv2` ($32 \to 64$ channels, $3 \times 3$ kernel, padding=1) each followed by ReLU activation and $2 \times 2$ max-pooling. The pooled feature map ($64 \times 16 \times 16$) is flattened and passed through a fully connected layer ($64 \cdot 16 \cdot 16 \to 128$ units) with ReLU, then through a final linear layer ($128 \to 1$) plus sigmoid to yield a probability of spot presence. At runtime, the GUI loads the pretrained weights from `spot_detection_cnn.pth`. During analysis, the user first selects a reference channel for spot detection using the tracking algorithms; for each detected spot, a fixed-size crop from the test channel is fed to the CNN, whose output probability is thresholded (user-adjustable) to produce a binary colocalization decision.
- **Intensity threshold approach**. This method uses a fluorescence intensity cutoff to determine whether the spots in the reference and test channels are colocalized. For each reference spot centroid, the algorithm checks whether the same coordinates (within a specified radius) in the test channel exhibit a local fluorescence intensity above the user-defined cutoff.
- **Manual detection approach**. Manual detection is implemented by showing the user side-by-side crops for the reference and test channels. The user can select a checkbox to determine the presence of colocalized spots. This method can be pre-populated with the results obtained from the machine learning and intensity threshold approaches, allowing the user to only manually verify a subset of samples.

#### Correlation Analyses

The `MicroLive` GUI enables users to perform autocorrelation analysis on the intensity time courses of individual fluorescent spots to quantify temporal persistence and fluctuation dynamics. The autocorrelation function (ACF) of a fluorescence trace $I(t)$ is defined as

$$G(\tau) = \frac{\langle \delta I(t) \delta I(t+\tau) \rangle}{\langle I(t) \rangle^2}, \tag{9}$$

where $\delta I(t) = I(t) - \langle I(t) \rangle$ and $\tau$ is a time lag. This function measures how fluctuations in intensity at a given time $t$ correlate with those at a later time $t + \tau$. Before computing autocorrelations, trajectories that are too short or have insufficient signal-to-noise can be filtered out to ensure reliable statistics. Additional noise-mitigation steps can be applied, for example, the autocorrelation at $\tau = 0$ (which can be artificially inflated by shot noise) can be replaced by an interpolated value from the first few non-zero lag points, and any baseline offset at long $\tau$ can be subtracted so that the correlation approaches zero at large lag. The resulting ACF curves (averaged over all trajectories detected in the ROI) can be fitted with linear or exponential decay models to extract the correlation decay time $\tau_c$. Bootstrap resampling of trajectories is used to estimate standard errors (6).

Biophysical parameters were estimated as in Larson et al. (7), that is, elongation rates were approximated as follows:

$$k_e \approx \frac{L}{\tau_c}\,, \tag{10}$$

where $L$ is the gene length, and initiation rates were approximated as follows:

$$k_i \approx \frac{1}{G(0) \cdot \tau_c}\,, \tag{11}$$

where $G(0)$ is the value of the autocorrelation function at $\tau = 0$.

### Verification Using a Synthetic Dataset

To assess the performance and accuracy of our `MicroLive` GUI, we compared its outputs with a known ground-truth synthetic dataset generated using the rSNAPed library, where all parameters are predetermined (8). In short, the rSNAPed library simulates live-cell single-molecule translation movies by first importing a real cell video loaded with fluorophores as the background of the movie; then adding simulated translation spots with fluctuating fluorescence intensities calculated using a TASEP (Totally Asymmetric Exclusion Process) model, which takes into account the ribosomal initiation rates, elongation rates, and ribosomal exclusion. The intensity of each spot is proportional to the number of simulated ribosomes and their elongation rates. This simulated movie provides an approach to verify the performance and accuracy of the GUI, as we know the true photobleaching decay rate and the true characteristics of the simulated spots, including spot size, spot intensity, number of spots, diffusion rate, and the initiation and elongation rates. We used rSNAPed to simulate the translation of smHA-KDM5B-BoxB-MS2 (1,910 codons, plasmid sequence is provided in the GitHub repository) using the parameters given in Table S1. We loaded the simulated movie into `MicroLive`, processed it, and calculated all parameters relevant to spot properties and dynamics. We found a strong agreement between the ground-truth dataset and `MicroLive` quantification.

| **Parameter** | **rSNAPed** | `Microlive` |
|---|---|---|
| $k_e$ | 5 aa/s | 5.2 aa/s |
| $k_i$ | 0.04 $s^{-1}$ | 0.038 $s^{-1}$ |
| Diffusion rate | 0.1 $px^2/s$ | 0.092 $px^2/s$ |
| $\sigma_{\text{psf}}$ | 1.5 px | 1.55 px |
| Photobleaching decay rate | 0.001 $s^{-1}$ | 0.001 $s^{-1}$ |

Table S1: Parameters used in rSNAPed for generating a simulated single-molecule translation movie of smHA-KDM5B-BoxB-MS2 construct.

#### Testing with a Real Microscopy Dataset

To evaluate the performance of `MicroLive` on real experimental data, we tested the GUI on our acquired microscopy single-molecule translation movies of smHA-KDM5B-BoxB-MS2 (9). First, we generated a U-2 OS cell line stably expressing both anti-HA-frankenbdoy-HaloTag and tdMCP-tdmSG probes as well as $\lambda$N-CAAX for tethering mRNAs to the plasma membrane upon binding the BoxB stem loops in the 3' UTR, all of which are under a Tet-On promoter. The day before imaging, we transiently transfected the stable cells seeded on MatTek chambers with the smHA-KDM5B-BoxB-MS2 construct in Opti-MEM for 3h, then replaced the medium with DMEM(+) containing 1 $\mu$g/mL doxycycline and 100 $\mu$M of JF646 dyes. On the imaging day, the cells were washed three times and fed with 1mL of DMEM(+) medium without phenol-red for imaging. We acquired time-lapse movies (600 frames at 1 fps, single $z$ plane, and a pixel size of 130 nm) on our Leica Stellaris 5 confocal microscope using a 63x oil immersion objective, 2% laser power at 488 nm (RNA channel) and 5% at 638 nm (nascent chain channel). The LIF files were imported into `Microlive`

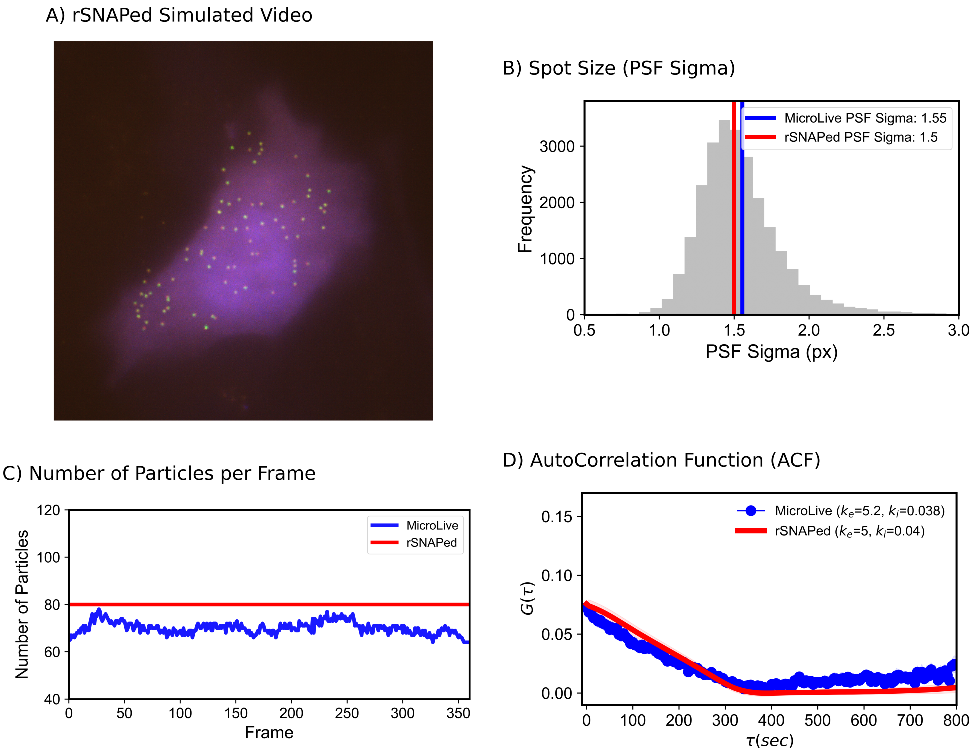


Figure S2: `Microlive` image processing results for a synthetic image generated by the rSNAPed library. A) The top image shows a simulated microscopy image, representing the translation of the smHA-KDM5B-BoxB-MS2 construct using the rSNAPed library and the parameters provided in Table S1. B) Comparison of the spot size used to generate the simulated dataset (red line) with the quantified spot size by `Microlive` (blue line). C) Time course showing the number of detected trajectories longer than 25 frames. 2D-tracking was performed using a particle size of 5 pixels and a search range of 7 pixels. The red line indicates the number of particles used for the simulation. D) Autocorrelation function calculated using the simulated data from rSNAPed (red line), and autocorrelation function calculated using the intensity recovered from tracked spots with `MicroLive` (blue line).

for cell segmentation, particle tracking, spot intensity extraction, and autocorrelation analysis. `Microlive` reliably detected spots and linked trajectories longer than 25 frames, yielding a mean decorrelation time of 122 seconds (Figure S3).

### Documentation

Technical and user documentation for `MicroLive` is provided in its GitHub repository `https://github.com/ningzhaoAnschutz/microlive` in the following links:

| **Document** | **GitHub Link** |
|---|---|
| Complete user manual | user_guide.md |
| Step-by-step tutorials | tutorial.md |
| Technical API documentation | api_reference.md |

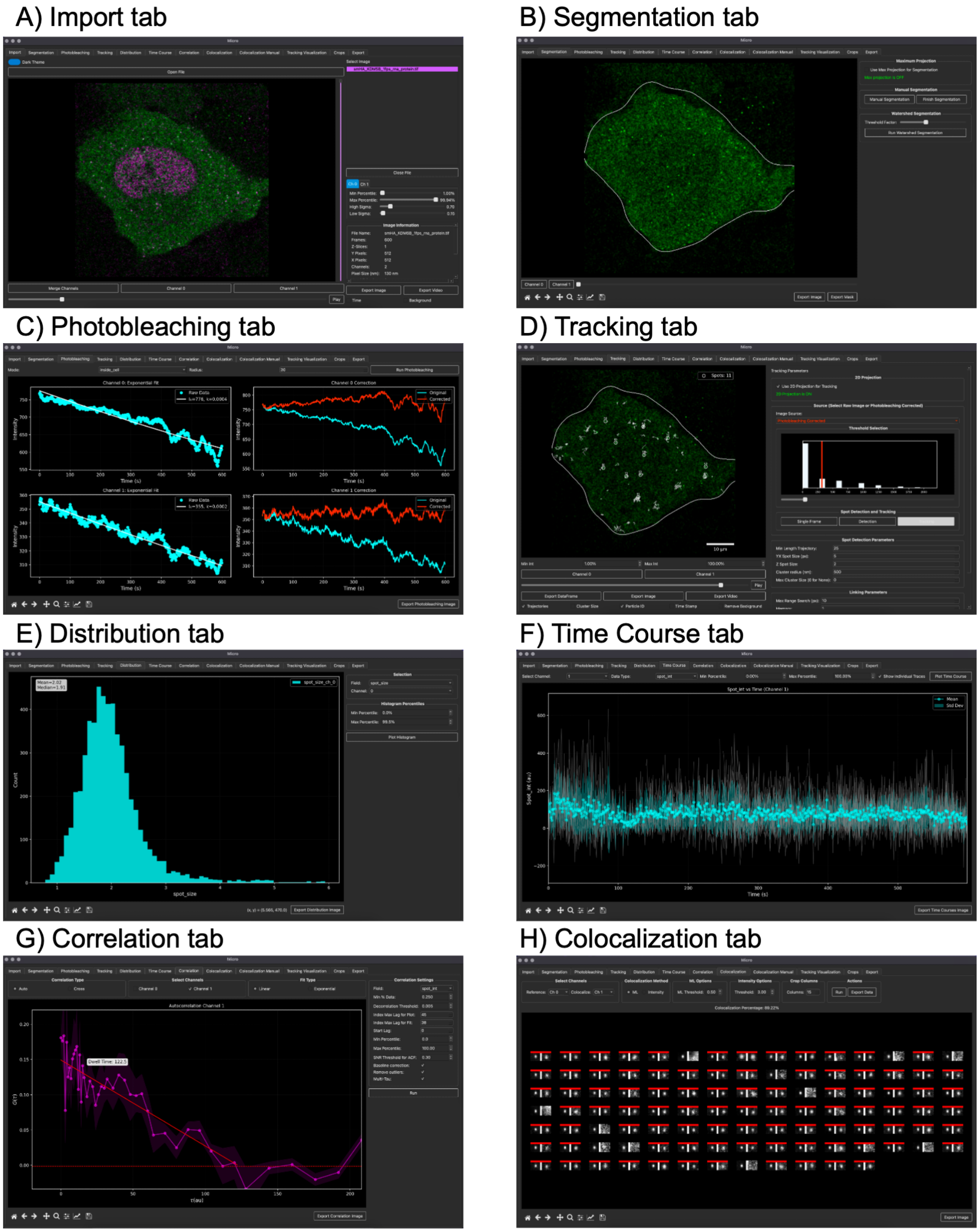


Figure S3: Screenshots show the main tabs in `Microlive`. A) Import tab, showing a representative microscopy movie where the green channel represents the mRNA channel, and the magenta channel represents the nascent chain channel. B) Segmentation tab. C) Photobleaching tab. D) Representative image showing the particle tracking results at a given time frame. E) Distribution tab showing the spot size for all spots detected in all frames. F) Time course tab showing the particle intensity over time. G) Correlation tab. A representative plot showing the autocorrelation function of intensity calculated from the trajectories of the nascent chains. H) Colocalization tab, showing pairs of crops for the mRNA channel and the protein channel. The red bar at the top indicates where the code is detecting a pair of colocalized spots.

### References


[1] Luc Vincent, and Pierre Soille. Watersheds in digital spaces: an efficient algorithm based on immersion simulations. *IEEE Transactions on Pattern Analysis & Machine Intelligence.* 13(06), pages 583-598. 1991.

[2] Kota Miura. Bleach correction ImageJ plugin for compensating the photobleaching of time-lapse sequences. *F1000Research.* 9, pages 1494. 2020.

[3] Kenneth Lyon, Luis U. Aguilera, Tatsuya Morisaki, Brian Munsky, and Timothy J. Stasevich. Live-cell single RNA imaging reveals bursts of translational frameshifting. *Molecular Cell.* 75(1), pages 172-183. 2019.

[4] Arthur Imbert, Wei Ouyang, Adham Safieddine, Emeline Coleno, Christophe Zimmer, Edouard Bertrand, Thomas Walter, and Florian Mueller. FISH-quant v2: a scalable and modular tool for smFISH image analysis. *RNA.* 28(6), pages 786-795. 2022.

[5] Daniel B. Allan, Thomas Caswell, Nathan C. Keim, Casper M. van der Wel, and Ruben W. Verweij. Soft-matter/trackpy: V0.7. *Zenodo* https://doi.org/10.5281/zenodo.16089574. 2025.

[6] Antoine Coulon, and David R. Larson. Fluctuation analysis: dissecting transcriptional kinetics with signal theory. *Methods in Enzymology.* Vol. 572, pages 159-191. 2016.

[7] Daniel R. Larson, Daniel Zenklusen, Bin Wu, Jeffrey A. Chao, and Robert H. Singer. Real-time observation of transcription initiation and elongation on an endogenous yeast gene. *Science.* 332(6028), pages 475-478. 2011.

[8] William S. Raymond, Sadaf Ghaffari, Luis U. Aguilera, Eric Ron, Tatsuya Morisaki, Zachary R. Fox, Michael P. May, Timothy J. Stasevich, and Brian Munsky. Using mechanistic models and machine learning to design single-color multiplexed nascent chain tracking experiments. *Frontiers in Cell and Developmental Biology.* 11, 1151318. 2023.

[9] Gabriel Galindo, Gretchen M. Fixen, Amelia Heredia, Tatsuya Morisaki, and Timothy J. Stasevich. All Probes Plasmids (APPs) for multicolor and long-term tracking of single-mRNA translation dynamics. *Molecular Biology of the Cell.* 36(6):mr6. 2025.
